# MCT4 and CD147 co-localize with MMP14 in invadopodia and autolysosomes and collectively stimulate breast cancer cell invasion by increasing extracellular matrix degradation

**DOI:** 10.1101/2023.09.03.556100

**Authors:** Signe Meng, Ester E. Sørensen, Muthulakshmi Ponniah, Jeppe Thorlacius-Ussing, Roxane Crouigneau, Magnus T. Borre, Nicholas Willumsen, Mette Flinck, Stine F. Pedersen

## Abstract

The lactate-proton cotransporter MCT4 and its chaperone CD147 are upregulated in breast cancers, correlating with decreased patient survival. Here, we test the hypothesis that MCT4 and CD147 favor breast cancer invasion through interdependent effects on extracellular matrix (ECM) degradation. MCT4 and CD147 expression and membrane localization were strongly reciprocally interdependent in MDA-MB-231 invasive breast cancer cells. Knockdown (KD) and overexpression (OE) of MCT4 and/or CD174 in- and decreased, respectively, migration, invasion, and fluorescent gelatin degradation. OE of both proteins increased gelatin degradation and appearance of the matrix metalloprotease (MMP)-generated collagen-I cleavage product reC1M more than each protein alone, suggesting a concerted role in ECM degradation. MCT4 and CD147 co-localized strongly with invadopodia markers at the plasma membrane and with MMP14, the lysosomal marker LAMP-1, and in some cases the autophagosome marker LC3, in F-actin-decorated, large intracellular vesicles.

We conclude that MCT4 and CD147 reciprocally regulate each other and support migration and invasiveness of MDA-MB-231 breast cancer cells in an interdependent manner. Mechanistically, this involves the MCT4-CD147-dependent stimulation of ECM degradation and specifically of MMP-mediated collagen-I degradation. We suggest that the MCT4-CD147 complex is co-delivered to invadopodia with MMP14.

## Introduction

Despite major advances in therapy in recent decades, breast cancer remains the leading cause of cancer mortality in women worldwide (Sung et al., 2021). Metastasis to bone, lungs, liver and brain account for about 90% of breast cancer deaths (Liang et al., 2020, Weigelt et al., 2005), highlighting the major unmet need for improved therapeutic strategies targeting mechanisms of metastasis in this disease. While the metastatic cascade through which breast cancer cells eventually colonize a distant organ and establish a metastasis involves multiple steps, a fundamental prerequisite is the ability of the cancer cells to remodel and degrade extracellular matrix (ECM), enabling them to invade through surrounding stroma and tissues (Mego et al., 2010, Lu et al., 2011).

The monocarboxylate transporter MCT4 (SLC16A3) is typically strongly expressed in highly glycolytic tissues, where it exports lactate and H^+^ derived from fermentative glycolysis, driven by the combined electrochemical gradients of these metabolites (Halestrap, 2013, Payen et al., 2020). The plasma membrane localization, and thus the transport function, of MCT4 are dependent on its chaperone CD147 (aka EMMPRIN or Basigin), a single pass transmembrane protein with multiple glycosylated states (Grass et al., 2014, Kirk et al., 2000). Conversely, CD147 expression was shown to be reduced upon knockdown of MCT4 (Gallagher et al., 2007, Andersen et al., 2018), indicating that the two proteins are interdependent. CD147 has also been assigned a role in binding of carbonic anhydrase IX to the CD147-MCT4 complex, facilitating the lactate-H^+^ transport (Ames et al., 2020).

Both MCT4 and CD147 are upregulated in many cancers, and their expression correlates with poor prognosis (Payen et al., 2020, Xin et al., 2016). In the case of MCT4, this is often attributed to its role in enabling the high fermentative glycolytic rates common in aggressive cancers, and hence enabling tumor growth (Morais-Santos et al., 2015, Le et al., 2011, Andersen et al., 2018). However, more recently, we and others have demonstrated roles for MCT4 in cancer cell migration and invasiveness (Gallagher et al., 2009, Kong et al., 2016, Zhu et al., 2014). Although a role for MCT4 interaction with integrins has been proposed to favor cell migration (Gallagher et al., 2009), the mechanisms through which MCT4 favors invasiveness remain essentially unelucidated.

It has long been appreciated that CD147 facilitates cancer invasiveness in a manner linked to production and activation of several matrix metalloproteinases (MMPs) (Grass et al., 2012, Sun and Hemler, 2001). However, the mechanisms involved have been contentious, including whether the ability of CD147 to regulate MMPs is independent from its role as an MCT chaperone (Schneiderhan et al., 2009). CD147 can reside in the plasma membrane, in subcellular compartments, and can be released from cells in various forms (Albrechtsen et al., 2019, Grass and Toole, 2015, Egawa et al., 2006, Taylor et al., 2002, Ko et al., 2023). The relation between CD147 glycosylation state, localization and function is incompletely understood, with both high-(HG) and low-glycosylated (LG) CD147 suggested to have the capacity for activating MMPs, although the HG form may do so more effectively (Huang et al., 2013, Bai et al., 2014). Finally, CD147 interacts with multiple other proteins with roles in cancer cell invasion, including CD44 (Grass et al., 2014), caveolin-1 and several integrins (Berditchevski et al., 1997, Tang and Hemler, 2004). Thus, whether the roles of MCT4 and CD147 in cancer cell motility are dependent on their interaction remains unknown.

The aim of the present study, therefore, was to test the hypothesis that MCT4 and CD147 favor breast cancer invasiveness through interdependent effects on ECM degradation. We show that MCT4 and CD147 expression and localization are highly interdependent, and that they co-localize not only in the plasma membrane but also in characteristic F-actin decorated vesicles. Overexpression of MCT4 and/or CD147 increases, and their knockdown decreases, invasiveness and ECM degradation. We suggest that the interaction of MCT4 and CD147 in both the plasma membrane and these vesicles is important for their role in breast cancer cell invasiveness.

## Results

### Protein expression levels of MCT4 and CD147 are reciprocally interdependent

The first step in understanding the possible relationship between MCT4 and CD147 in regulation of breast cancer cell motility was to determine their interrelationship at the protein level. Western blotting (WB) showed that MCT4 protein expression was essentially ablated 48 h after transfection of MDA-MB-231 human breast cancer cells with MCT4 siRNA (Fig. 1a-bi). Notably, this MCT4 knockdown (KD) was associated with a strong reduction in the expression of high-glycosylated (HG) CD147 (Fig. 1a-bii). Conversely, KD of CD147 using two different siRNAs not only strongly attenuated CD147 expression but also essentially ablated MCT4 expression (Fig. 1c-d). A different pattern emerged when MCT4 or CD147 were instead transiently overexpressed (OE). OE of MCT4, but not of CD147, significantly increased the MCT4 protein level (Fig. 1e-fi). Similarly, only OE of CD147 increased the protein level of CD147, and this was only significant for the LG form (Fig. 1e-fii-iii). Expression levels of both proteins were the same upon their co-expression using the same total amount of cDNA, i.e. 50% of each, however, a misfolded MCT4 dimer band (Wilson et al., 2009) disappeared upon co-expression (Fig. 1e-f). To further interrogate an apparent tendency for an increase in MCT4 level upon CD147 expression (Fig. 1fi), we determined the mRNA levels of *SLC16A3* and *BSG* by qPCR analysis (Fig. 1g-h). The pattern of changes was similar to that seen at the protein level, but, interestingly, revealed a minor but significant increase in MCT4 mRNA level upon CD147 expression (Fig. 1g).

**Figure 1.**
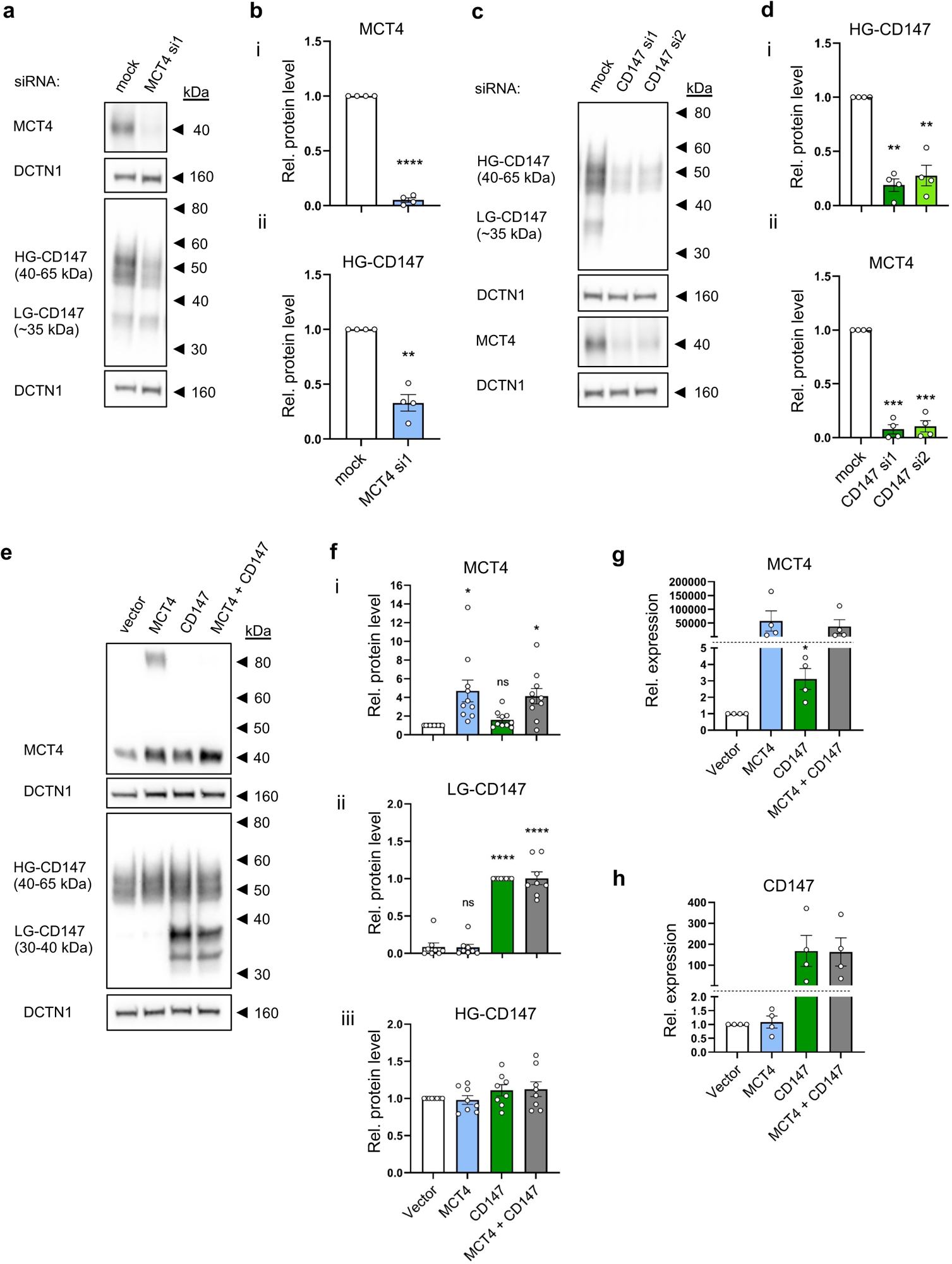
Protein expression levels of MCT4 and CD147 are reciprocally interdependent. MDA-MB-231 cells were transfected with siRNAs targeting MCT4 (**a, b**), CD147 (**c, d**) or scrambled siRNA control (mock), or with MCT4, CD147 or MCT4 and CD147 plasmids or empty pcDNA3.1 (+) vector as negative control (**e-h**). 48 h (KD) or 24 h (overexpression) post-transfection, cells were lysed and subjected to Western blot or qPCR analysis as indicated. **(a-d)** Representative Western blots and corresponding quantification of MCT4 and CD147 protein levels. DCTN1 was used as loading control. MCT4 and HG-CD147 band intensities were normalized to their respective loading controls and control (mock). n=4. **(e-f)** Representative Western blots and corresponding quantification of MCT4 and CD147. DCTN1 was used as loading control. MCT4 and CD147 band intensities were normalized to loading controls and control (vector). n=10 (MCT4), 8 (CD147). **(g-h)** Relative mRNA levels (qPCR) of MCT4 and CD147. Data was normalized to housekeeping genes β-actin and TBP and to vector control. n=4. Statistical analysis: Two-tailed paired Student’s *t*-test (b, g), one-way ANOVA with Dunnett’s post-test (d, f, h). *, **, ***, ****: p < 0.05, 0.01, 0.001, 0.0001.

We speculated that the apparent lack of MCT4-CD147 interdependence after overexpression could reflect the high basal expression of both proteins in MDA-MB-231 cells and repeated the experiments in MCF10A non-cancer mammary epithelial cells and MCF-7 breast cancer cells, a non-invasive luminal breast cancer cell type (Suppl. Fig. 1). In both cell types, endogenous levels of especially MCT4 were much lower than in MDA-MB-231 cells, and while CD147 expression alone did not increase MCT4 expression, co-expression with CD147 resulted in much higher levels of MCT4 than expression of MCT4 alone, and strongly reduced the misfolded dimer band (Suppl. Fig. 1a). In both cell types, co-expression furthermore appeared to increase levels of the fully glycosylated HG form of CD147. This pattern was confirmed by quantification of multiple such experiments in MCF10A cells (Suppl. Fig. 1b-d).

Taken together, these results show that protein levels, and most likely also the correct folding and maturation, of MCT4 and CD147 are reciprocally interdependent.

### Overexpression increases MCT4 and CD147 plasma membrane localization and cellular lactate extrusion

We next asked whether OE of MCT4 and/or CD147 led to upregulation of functional, plasma membrane-localized proteins. We overexpressed the two proteins alone or together as before, followed by immunofluorescence microscopy (IFM) analysis of total levels of both proteins (endogenous + OE) (Fig. 2a). MCT4 (magenta) and CD147 (green) are visible both in intracellular structures and in the plasma membrane (arrowheads), where they partially co-localize (white) both endogenously and after OE. To quantify membrane localization, we first performed blinded line scan analysis of the IFM images (Materials and Methods; Suppl. Fig. 2). Determined in this manner, OE of MCT4 and/or CD147 increased, or tended to increase, plasma membrane localization of both proteins (Fig. 2c-d). As seen in the zoomed IFM images (Fig. 2a) and illustrated by representative line scan profiles (Fig. 2b), OE was also associated with increased intracellular levels of the protein. This was particularly prominent for MCT4 alone, consistent with the misfolded MCT4 dimer seen by WB (Fig. 1, Suppl. Fig. 1), but was also detectable for CD147 alone and for the co-expressed proteins, consistent with the high level of LG-CD147 observed in Fig. 1.

**Figure 2.**
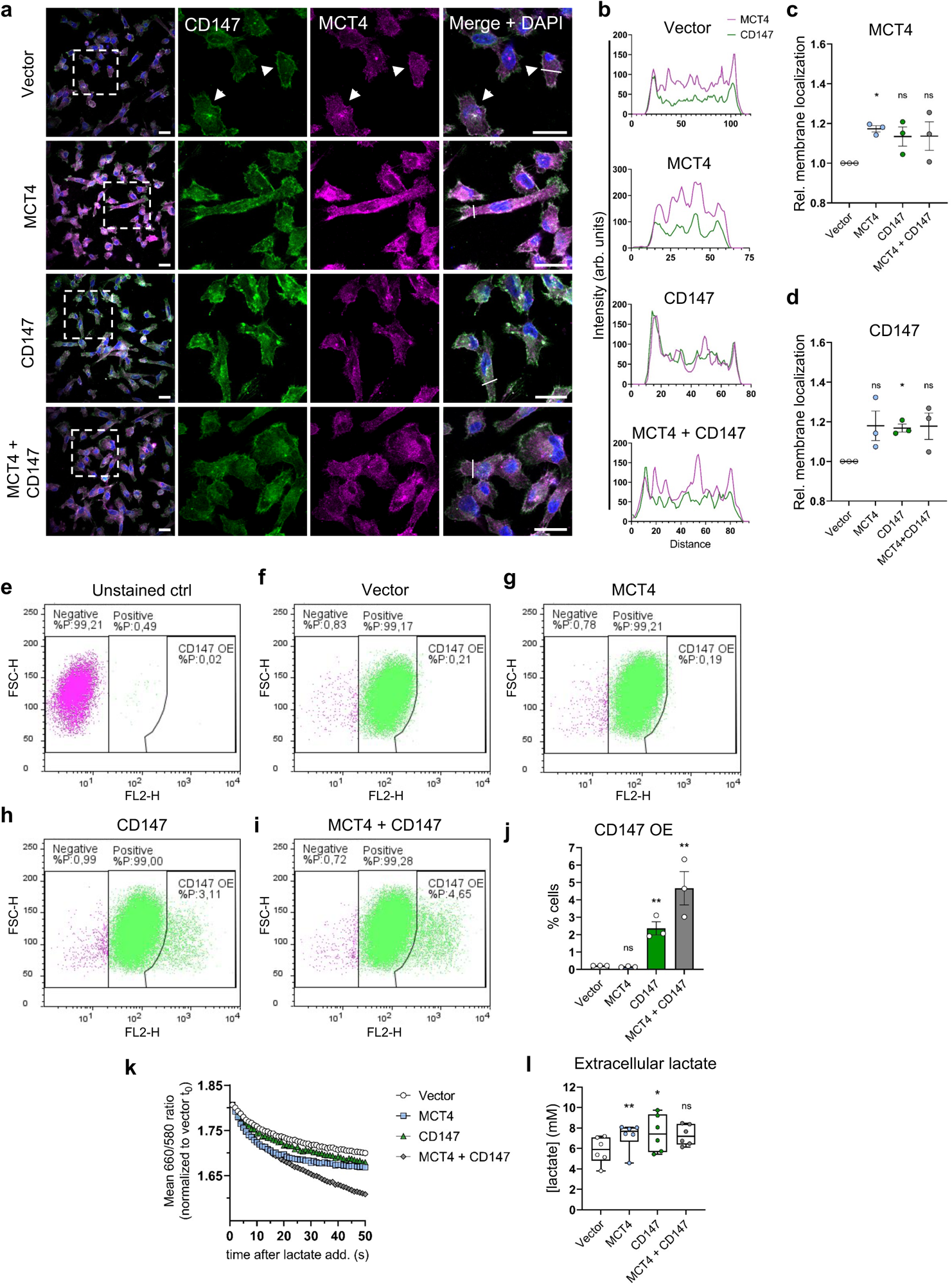
Overexpression increases MCT4 and CD147 plasma membrane localization and cellular lactate extrusion. **(a)** Representative overview images (left) and zooms (right) of MDA-MB-231 cells transfected with empty vector, MCT4, CD147 or MCT4 and CD147 plasmids. Cells were grown for 24 h, fixed and subjected to IFM analysis with primary antibodies targeting MCT4 (magenta) and CD147 (green). Nuclei were stained with DAPI (blue). White arrowheads show co-localization of MCT4 and CD147 in the plasma membrane and white lines represent the area analyzed by line scan analysis in (b). Scale bars: 20 µm. n=3. **(b)** Representative line scan analysis plots showing intensities (MCT4 magenta, CD147 green) along a selected line (white lines in (a)). **(c-d)** Relative membrane localization of MCT4 and CD147, determined by line scan analysis. Color intensities were normalized to cell count and to their respective control (vector). n=3. **(e-i)** Representative flow cytometric dot plots (FSC vs. FL2 (PE anti-human CD147). n=3 **(j)** Quantification of CD147 overexpression population. Statistical analysis was performed on log-transformed data. n=3. **(k)** Relative change in pH_i_ of MDA-MB-231 cells transfected with MCT4, CD147, or both plasmids, or with empty pcDNA3.1 (+) vector as negative control, measured as the change in SNARF-AM ratio after addition of 20 mM Na^+^-lactate. Representative experiment showing the mean 660/580 nm ratio for 50 s after adding 20 mM lactate. The data was normalized to vector t_0_. **(l)** Lactate concentration (mM) in culture medium from MDA-MB-231 cells cultured for 48 h after transfection with MCT4, CD147 or MCT4 and CD147 plasmids or empty pcDNA3.1 (+) vector as negative control. n=6. Statistics: (c), (d), (j), (l): one-way ANOVA followed by Dunnett’s multiple comparisons post-test.

Flow cytometric analysis of the plasma membrane level of CD147 using an independent, directly PE-conjugated antibody confirmed its increased membrane localization upon CD147 OE and suggested that co-expression with MCT4 might further increase CD147 membrane localization (Fig. 2e-j). Because MCTs co-transports lactate and H^+^ bidirectionally driven only by the electrochemical gradients of these ions (Halestrap, 2013), and as MDA-MB-231 cells do not express MCT1 (Asada et al., 2003, Andersen et al., 2018), MCT4 activity can be measured as the rate of change in intracellular pH (pH_i_) upon addition of lactate. To determine whether MCT4 OE increased lactate transport capacity, we therefore first loaded MDA-MB-231 cells with SNARF-AM and determined the intracellular acidification rate upon addition of 20 mM Na^+^-lactate to the Ringer solution (Fig. 2k). The initial rate and the duration of intracellular acidification appeared to be increased by OE of MCT4 or both proteins together, but effects were modest and not easily quantified. We therefore additionally determined lactate levels of culture medium collected 48 h after transfection, which were significantly increased after OE of either MCT4 or CD147 (Fig. 2l).

These results show that MCT4 and CD147 localize both intracellularly and to the plasma membrane, partially colocalizing in both locations, and that plasma membrane localization of both, as well as cellular lactate extrusion, are increased upon their overexpression.

### Invasion of MDA-MB-231 cells is dependent on MCT4 and CD147

Both MCT4 (Kong et al., 2016, Zhu et al., 2014) and CD147 (Grass et al., 2012, Sun and Hemler, 2001) have been implicated in cancer cell invasion, however, their interdependence in this regard has not been addressed, and the underlying mechanisms are unclear. To gain insight into this, we knocked down and overexpressed both proteins, alone and in combination, and studied the effects on MDA-MB-231 cell motility.

Transient siRNA mediated KD of MCT4 (which reduced MCT4 protein level by ∼90% and HG-CD147 protein level by ∼70%, see Fig. 1b) reduced MDA-MB-231 cell migration and invasion 24 h after seeding in Boyden chamber assays by ∼50% (Fig. 3a-c). Confirming this result, stable shRNA-mediated KD of MCT4, which reduced MCT4 and HG-CD147 protein levels by ∼90% and ∼70%, respectively (Suppl. Fig. 3), similarly reduced cell migration and invasion (Fig. 3b-c). This effect was not due to an impact of MCT4 KD on cell proliferation, which, although decreased after transient KD of MCT4 was unaltered after its stable KD (Suppl. Fig. 4a-b). When we instead knocked down CD147 (reducing HG-CD147 and MCT4 protein levels by ∼80 and ∼90%, respectively, Fig. 1d), a similar reduction in migration and invasion was observed, using two different siRNAs (Fig. 3d-f) and without effect on proliferation (Suppl. Fig. 4c). Finally, when MCT4, CD147, or both, were overexpressed in MDA-MB-231 cells, and migration and invasion determined 24 h after seeding, invasion was increased about four-fold (albeit not significant in CD147-transfected cells due to substantial variation), whereas migration was, interestingly, not significantly increased. This, again, was not due to effects on proliferation, which was unaffected in the overexpression conditions (Suppl. Fig. 4d).

**Figure 3.**
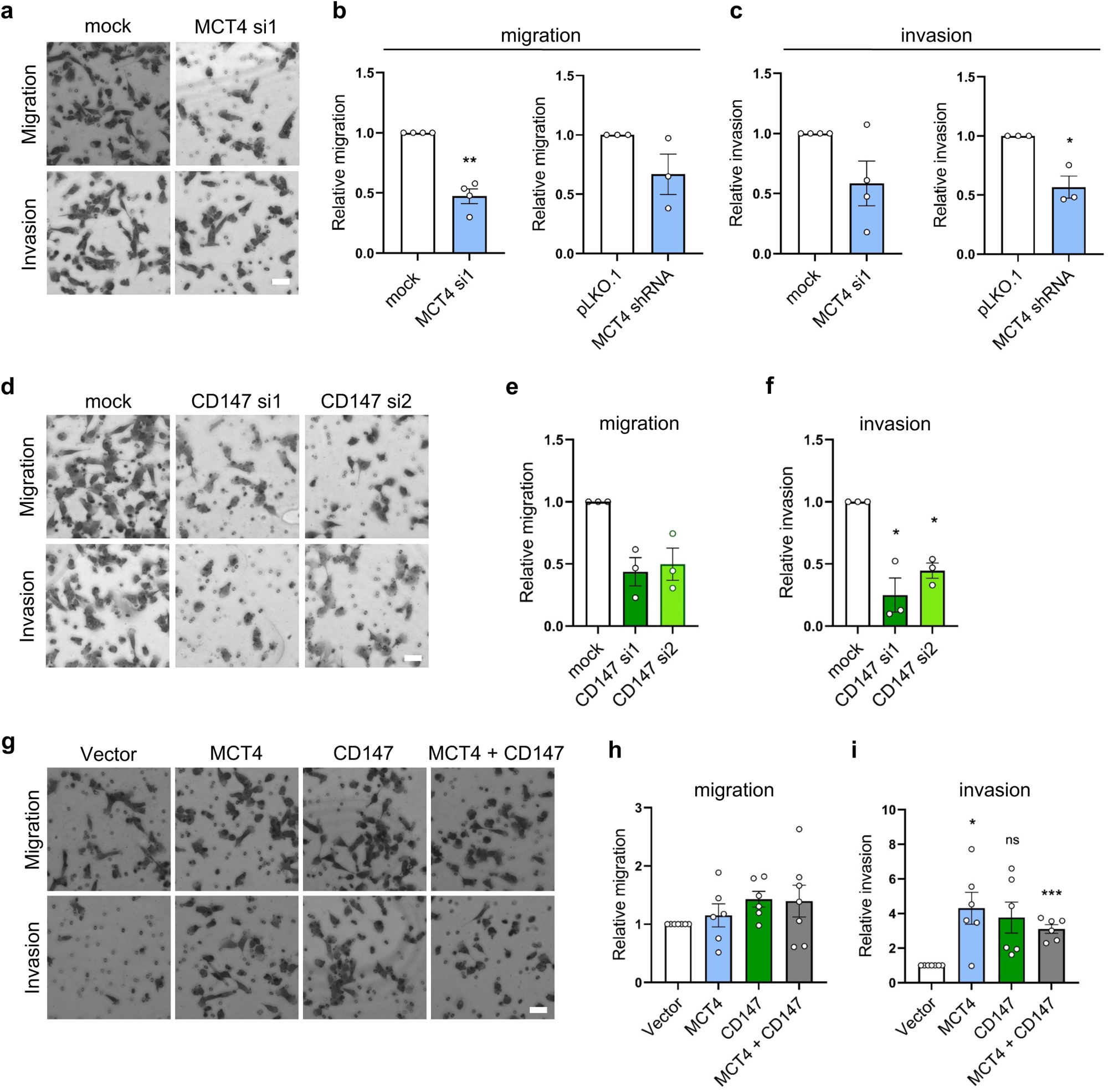
MCT4 and CD147 are both important for the invasiveness of MDA-MB-231 cells. MDA-MB-231 cells were transfected with siRNA targeting MCT4 (a-c) or CD147 (d-f) or with MCT4, CD147, both constructs or the corresponding empty vector (g-i). Alternatively, MDA-MB-231 cells with stable KD of MCT4 were employed (b-c). Cells were seeded in migration or invasion Boyden chamber inserts with 10% FBS as chemoattractant in lower chamber, and migration or invasion, as indicated, was determined 24 h after seeding of the cells in the upper chamber. Cells were fixed and stained with Giemsa solution and relative cell migration or invasion was determined by counting the cells which had moved through the inserts. **(a, d, g)** Representative images. Scale bar: 40 µm. **(b, e, h)** Relative cell migration as determined by the number of migrated cells relative to that in the control condition (mock, pLKO.1 or vector). **(c, f, i)** Relative cell invasion as determined by the number of cells invaded relative to that in the control condition (mock, pLKO.1 or vector). (a-i) n = 4, 3, 3, 6 in MCT4 siRNA KD, MCT4 stable KD, CD147 KD and overexpression, respectively. Statistics: Two-tailed, paired Students *t*-test (b, c) or one-way ANOVA with Dunnett’s post-test (e, f, h, i).

These results show that MCT4 and CD147 are necessary for MDA-MB-231 cell migration and invasion, and that their overexpression further increases invasion.

### MCT4 and CD147 are important for extracellular matrix degradation

The ability of the cancer cells to degrade the surrounding ECM is an essential element of their invasiveness. ECM degradation has previously been shown to be regulated by other net acid extruding transporters, at least in part because of the acidic pH optimum of MMPs (Busco et al., 2010, Pedraz-Cuesta et al., 2016). Given the role of MCT4 in coupled extrusion of lactate and H^+^, we therefore reasoned that the transporter could support invasiveness by facilitating ECM degradation. To test this hypothesis, we established a matrix degradation assay based on the ability of the cells to degrade Oregon-green conjugated gelatin, a collagen derivative with structural and functional properties similar to those of collagens (Bello et al., 2020). MDA-MB-231 cells were seeded on the fluorescent gelatin, and live imaging experiments established that cells actively degrade the matrix under them, as they move on the coverslip (Suppl. Video 1-2). Subsequently, cells were transfected with the relevant siRNAs or overexpression plasmids, fixed after 4, 8 or 12 h, and stained with Rhodamine-phalloidin and DAPI to visualize F-actin and nuclei (zoomed images in Fig. 4; full overview images in Suppl. Fig. 5 and Suppl. Fig. 6). Mock-transfected cells efficiently degraded the gelatin, evident as the gradual disappearance of Oregon-green fluorescence over time, particularly under actin-rich cellular structures, both protrusions and apparently intracellular regions (Fig. 4a, white arrowheads). In contrast, after MCT4 KD, cells appeared to degrade substantially less gelatin. Quantification of the degraded areas and normalization to cell count showed that gelatin degradation at time 8 h was reduced by more than 80% by MCT4 KD (Fig. 4b). CD147 KD similarly decreased gelatin degradation under actin-rich structures (Fig. 4c, white arrowheads), and reduced gelatin degradation at 8 h, although only by about 50% (Fig. 4c-d). MCT4 or CD147 KD also tended to reduce degradation at 12 h after seeding, although extensive experimental variability at this time precluded statistical significance (Fig. 4a-d). In contrast, overexpression of MCT4 or CD147 alone, or both proteins together, did not alter gelatin degradation, although a tendency for an increase was seen at 12 h upon combined overexpression (Fig. 4e-f).

**Figure 4.**
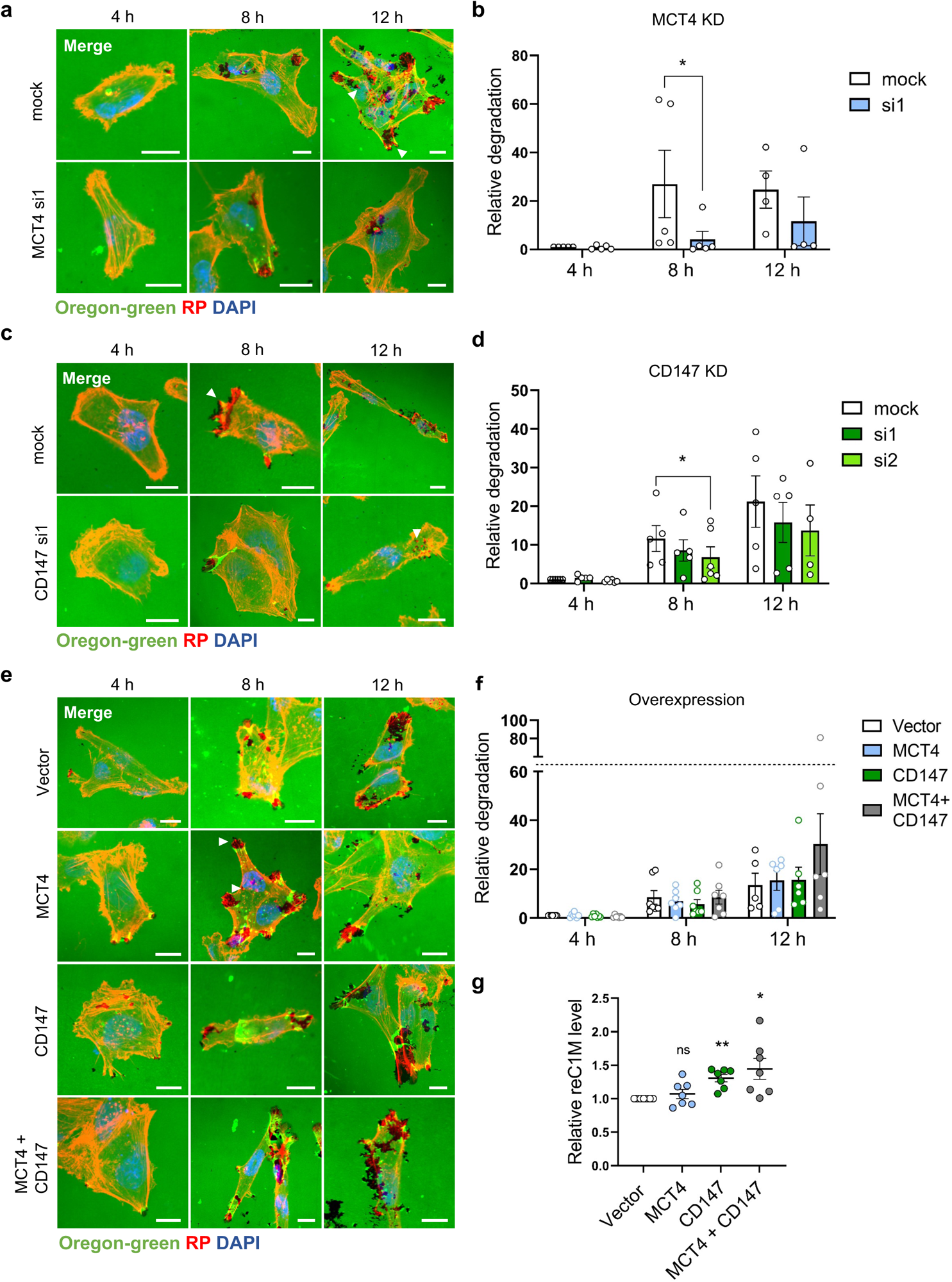
MCT4 and CD147 are important for extracellular matrix degradation. MDA-MB-231 cells were transfected with siRNA targeting MCT4 (a-b) or CD147 (c-d) or with empty vector, or corresponding MCT4-, CD147-, or MCT4- and CD147 plasmids (e-f), seeded on Oregon-green conjugated gelatin-coated coverslips and grown for 4, 8 or 12 h. Cells were subsequently fixed and stained for F-actin (Rhodamine Phalloidin, RP), and nuclei (DAPI). Suppl. Fig. 5 and 6 show the corresponding full overview images of the cropped images in this figure, as well as the images for CD147 siRNA-2. **(a, c, e)** Representative images of cells growing on Oregon-green gelatin for the indicated times, presented as merged images of Oregon-green, F-actin (RP) and DAPI. White arrowheads indicate overlap of gelatin degradation with F-actin staining at cell protrusions and intracellular structures. Scale bars: 10 µm. **(b, d, f)** Quantification of gelatin degradation (performed in ImageJ) by MDA-MB-231 cells after 4, 8, and 12 h. Relative degradation is the degraded area fraction normalized to the cell count. The data points represent the mean relative degradation of 6-13 images per condition. n= 4-5, 4-6, 5-7 for MCT4 KD, CD147 KD and overexpression, respectively. Statistics: (b), (d), (f): Two-tailed paired Students *t*-test (b) or one-way ANOVA followed by Dunnett’s multiple comparisons post-test (d, f) for each time point. Statistical analysis was performed on log-transformed data. **(g)** Relative reC1M (MMP-generated fragment of collagen-I) concentration measured by ELISA in samples collected from the culture medium of MDA-MB-231 cells (transfected with vector, MCT4, CD147 or MCT4 and CD147 plasmid constructs) 48 h after seeding on collagen-I. reC1M concentrations are normalized to the vector control within each independent experiment. n=7. One-way ANOVA followed by Dunnett’s multiple comparisons post-test was performed on log-transformed data.

These experiments strongly indicated that collagen degradation is dependent on MCT4 and CD147. To confirm this and gain insight into the nature of the degrading protease, we next overexpressed MCT4, CD147 or both in MDA-MB-231 cells as above, seeded the cells on collagen-I, and determined the appearance of reC1M, a collagen-I neoepitope generated specifically through its degradation by MMP-2, 9, or -13 (Leeming et al., 2011), in the culture medium by ELISA, 48 h after seeding (Fig. 4g). Notably, reC1M levels were significantly increased upon overexpression of CD147 or both proteins together.

These results show that MCT4 and CD147 are important for the degradation of ECM by MDA-MB-231 cells, and that this involves MMP-dependent degradation of collagen-I.

### MCT4 and CD147 co-localize to ECM-degrading invadopodia and MMP14-containing intracellular structures

Having shown that MCT4 and CD147 were important for matrix degradation by MDA-MB-231 cells, we wished to determine whether the two proteins co-localized in actively degrading invadopodia. To increase formation of invadopodia, we overexpressed constitutively active Src (Src Y527F) (Seals et al., 2005, Artym et al., 2006). Co-staining for Src, F-actin, and pY421-cortactin, which localizes to invadopodia and is important for their function (Jeannot and Besson, 2020) confirmed the strong localization of pY421-cortactin to invadopodia (Suppl. Fig. 7a). We could now use pY421-cortactin as an invadopodia marker, and its subsequent co-staining with MCT4 and CD147 showed that both proteins prominently, albeit only partially, co-localize to these structures (Suppl. Fig. 7b, white arrows).

To determine whether endogenous MCT4 and CD147 localized to actively degrading invadopodia in non-Src transformed cells, cells were cultured on Oregon-green gelatin for 8 h and stained for MCT4 or CD147, Oregon-green, and pY421-cortactin (Fig. 5a) or F-actin (Fig. 5b). This revealed clear localization of both proteins to pY421-cortactin- and F-actin rich structures, colocalizing with marked matrix degradation (white arrowheads). While some of these structures were reminiscent of lamellipodia or other focal adhesion-rich protrusions, others were intracellular, either vesicular or found at the interface between cells and matrix (Fig. 5b, lower row, arrowhead). Co-immunofluorescence analysis with F-actin, LAMP1 (lysosomes), protein disulfide-isomerase (PDI, endoplasmic reticulum (ER)), Golgin97 (Golgi apparatus) and TOM20 (mitochondria) indicated that of these organelles, CD147 and MCT4 were mainly associated with F-actin containing structures and with lysosomes, arguing against major accumulation in ER or Golgi apparatus (Suppl. Fig. 8).

**Figure 5.**
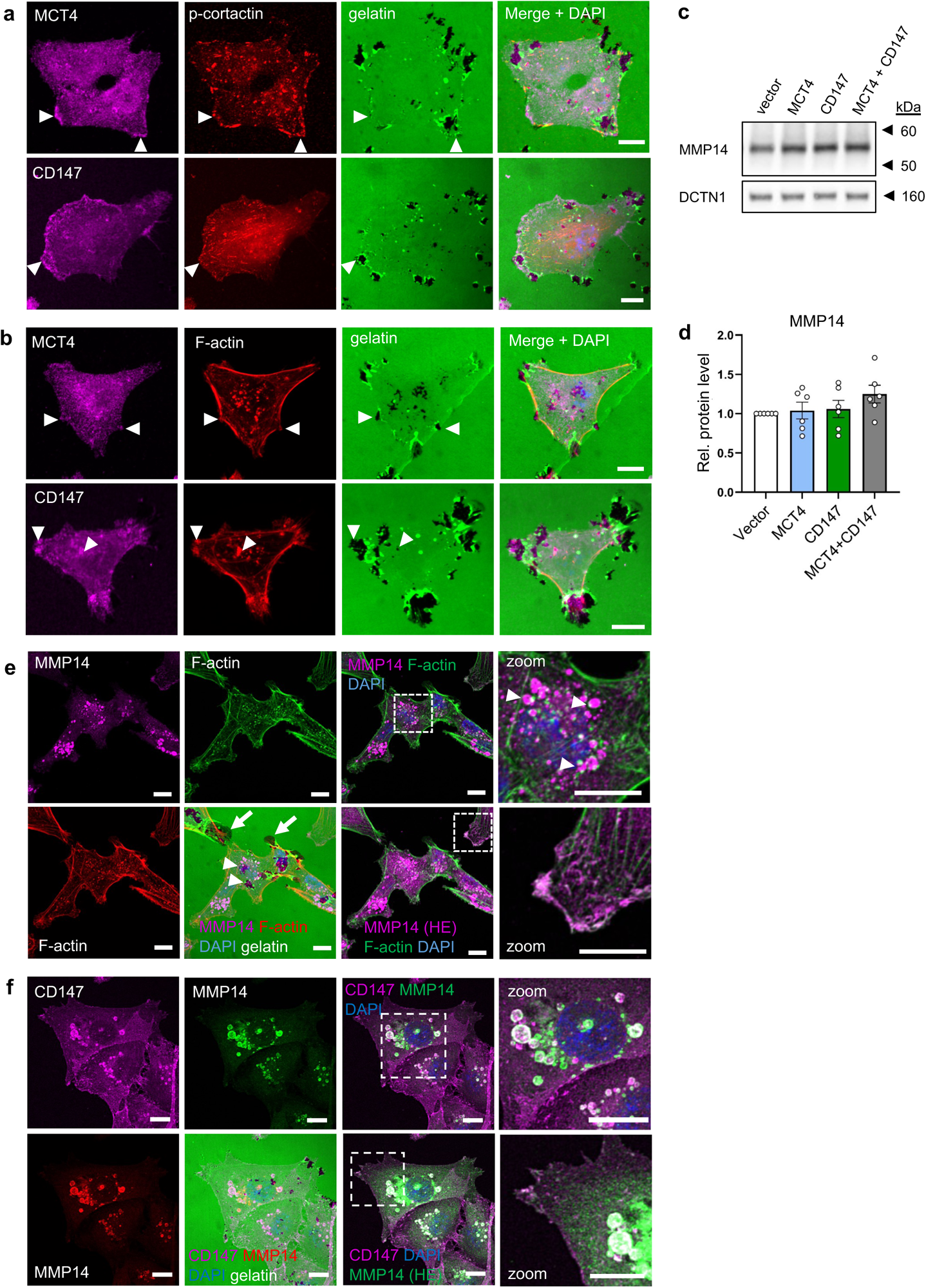
MCT4 and CD147 co-localize to ECM-degrading invadopodia and MMP14-containing intracellular structures. MDA-MB-231 wt cells were seeded on oregon-green conjugated gelatin-coated coverslips, fixed after 8 h of growth and subsequently subjected to IFM analysis. Nuclei were stained with DAPI (blue). **(a, b)** Representative images of MDA-MB-231 cells on gelatin (green) showing the cellular localization of MCT4 (magenta) or CD147 (magenta), and pY421-cortactin (red, in a) or F-actin (red in b). White arrowheads indicate overlap of gelatin degradation with MCT4/CD147 and pY421-cortactin (a) or F-actin (b). Scale bar: 10 µm. n=3. **(c,d)** Representative Western blot (c) and corresponding quantification (d) of MMP14. DCTN1 was used as loading control. MMP14 band intensities were normalized to loading control and control (vector). n=6. **(e)** Representative images of MDA-MB-231 cells on gelatin (green, lower panel) showing the cellular localization of MMP14 (magenta) and F-actin (green or red). MMP14 is shown at a low (upper panel) and high (HE, lower panel) exposure. Scale bar: 10 µm. n=3. **(f)** Representative images of MDA-MB-231 cells on gelatin (green, lower panel) showing the cellular localization of CD147 (magenta) and MMP14 (red or green). MMP14 (green) is shown at a low (upper panel) and high (HE, lower panel) exposure. Scale bar: 10 µm. n=3.

MMP14 is overexpressed in many cancers and is a major upstream activator of other MMPs. These include MMP2 (Itoh et al., 2001) and MMP13 (Knäuper et al., 2002), two of the MMPs generating the reC1M neoantigen (Leeming et al., 2011). Total MMP14 expression was not significantly altered by overexpression of MCT4, CD147, or both proteins (Fig. 5c-d). Growing the MDA-MB-231 cells on Oregon-green gelatin followed by staining for MMP14 and F-actin revealed that although also expressed at the plasma membrane (Fig. 5e, high exposure), MMP14 was most strongly localized to intracellular vesicles that each appeared characteristically coupled to F-actin “dots” (white arrowheads). Co-staining of MMP14 with CD147 on Oregon-green gelatin revealed clear co-localization between MMP14 and CD147 in these intracellular vesicles, and higher exposure also revealed co-localization at the plasma membrane (Fig. 5f). Matrix degradation was occasionally detectable associated with these structures (arrowheads) but was more frequently seen at the cell protrusions (arrows).

These results show that CD147 and MMP14 co-localize to matrix-degrading protrusions as well as to large intracellular vesicles decorated by F-actin “dots”. By comparison with the CD147-MCT4 (co-)localization to these structures (Fig. 2a, Fig. 5a, see also Suppl. Fig. 8) it can be inferred that these structures also contain MCT4.

### MMP14 and CD147 co-localize with MMP14 in LC3-positive vesicles

MMP14 in MDA-MB-231 cells is mainly localized to late endosomes/lysosomes from where it is trafficked to invadopodia (Monteiro et al., 2013, Poincloux et al., 2009). The striking F-actin “dots” on the MMP14- and CD147 vesicles were reminiscent of the F-actin comet-bearing vesicles formed during phagocytosis, macropinocytosis (Mylvaganam et al., 2021) and lysosome-autophagosome fusion (Nakamura and Yoshimori, 2017, Kast and Dominguez, 2017). Furthermore, MMP14 is known to play a critical role in collagen-I phagocytosis and (Lee et al., 2006). Co-immunofluorescence analysis for MMP14 and LAMP1 showed substantial staining overlap (Fig. 6a, bottom right, arrowheads) confirming the localisation of MMP14 to late endosomes/lysosomes. Interestingly, some gelatin degradation overlapped with these MMP14-LAMP1 organelles (Fig. 6a, top right, arrowhead). A fraction (arrowheads) of the vesicles (here stained by CD147) were also positive for microtubule associated protein 1 light-chain 3 (LC3) (Fig. 6b), suggesting that they could be autophagosomes, although other interpretations are possible (Runwal et al., 2019). Also MCT4 co-localized with LC3 in intracellular vesicles (Fig. 6c, top, arrowheads), whereas co-localisation of LC3 with LAMP1 appeared to be minimal (Fig. 6, lower panel).

**Figure 6.**
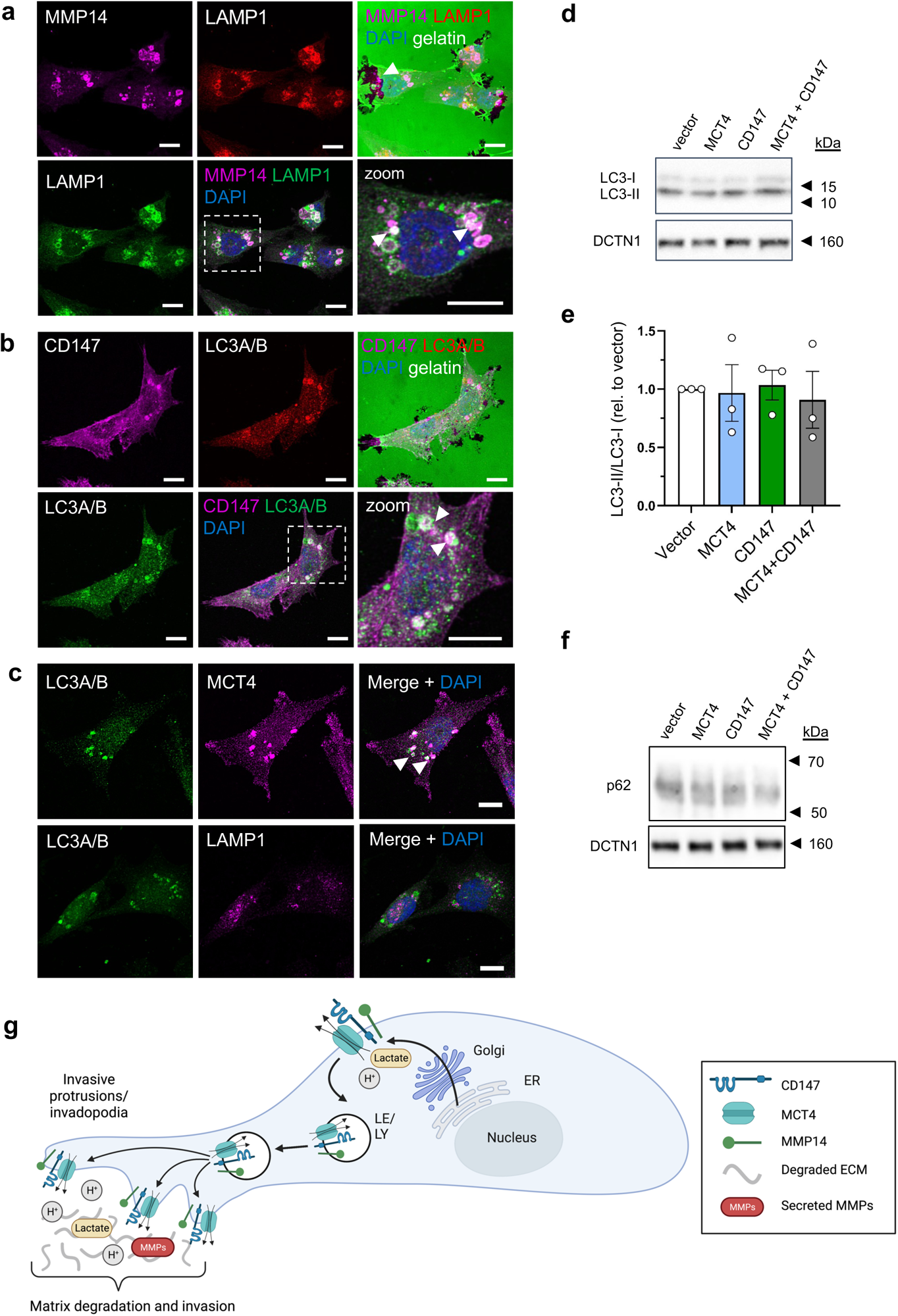
MCT4 and CD147 co-localize with MMP14 in LC3-positive vesicles. MDA-MB-231 wt cells were seeded on Oregon-green conjugated gelatin-coated coverslips, fixed after 8 h of growth and subsequently subjected to IFM analysis. Nuclei were stained with DAPI (blue). **(a)** Representative images of MDA-MB-231 cells on gelatin (green, upper panel) showing the cellular localization of MMP14 (magenta) and LAMP1 (red in upper panel, green in lower panel). Scale bar: 10 µm. n=3. **(b)** Representative images of MDA-MB-231 cells on gelatin (green, upper panel) showing the cellular localization of CD147 (magenta) and LC3 A/B (red in upper, green in lower panel). Scale bar: 10 µm. n=3. **(c)**. Representative images showing the localization of LC3 A/B (green) and MCT4 (magenta, top panel) and of LC3 A/B (green) and LAMP1 (magenta, lower panel). Scale bar: 10 µm. n=1. **(d-e)** Representative Western blot and corresponding quantification of LC3-II/I ratio 48 h after transfection with MCT4, CD147 or MCT4 and CD147 plasmids or empty pcDNA3.1 (+) vector as negative control. DCTN1 was used as loading control. n=3. **(f)** Representative Western blot of p62 protein level 48 h after transfection with MCT4, CD147 or MCT4 and CD147 plasmids or empty pcDNA3.1 (+) vector as negative control. DCTN1 was used as loading control. n=3. (g) Working hypothesis. MCT4-CD147-costructures are delivered to the plasma membrane along with MMP14 and from there the three proteins are retrieved together by endocytosis and delivered to invadopodia in large vesicles of late endosomal/lysosomal origin. Whether the role of MCT4-CD147 in ECM degradation involves a role in formation or recycling of the large vesicles and/or lactate-H^+^ efflux in invadopodia remains to be determined. See text for further details.

An increased ratio of the lipidated, autophagosome-associated form of LC3 (LC3-II) to LC3-I and a decrease in the protein level of the SQSTM1/p62 autophagy receptor (which is degraded during autophagy of its recruited targets) is indicative of increased autophagy. If MCT4 and CD147 played a role in the phagocytosis and intracellular degradation of collagen-I in such structures, and this represented a sizable fraction of total cellular autophagy, their overexpression could increase total cellular autophagic flux. However, neither LC3-II/LC3-I ratio or the protein level of the SQSTM1/p62 autophagy receptor were altered by overexpression of MCT4, CD147, or both proteins together (Fig. 6e-f).

These results show that MCT4, CD147 and MMP14 co-localize with LC3 in intracellular vesicles, but that their overexpression does not detectably impact total cellular autophagy.

## Discussion

Increased expression of MCT4 or CD147 correlates with poor prognosis in many cancers (Payen et al., 2020, Xin et al., 2016), and previous work from us and others has established roles for both MCT4 (Kong et al., 2016, Zhu et al., 2014) and CD147 (Grass et al., 2012, Sun and Hemler, 2001) in cancer cell migration and invasion. However, the mechanisms remain essentially unelucidated, and the interdependency of the two proteins for proper expression and localization (Gallagher et al., 2007, Andersen et al., 2018, Morais-Santos et al., 2015) makes it difficult to interpret such findings unless both proteins were accounted for in study designs. Here, we tested the hypothesis that MCT4 and CD147 favor breast cancer cell invasiveness through interdependent effects on ECM degradation.

Consistent with previous reports (Gallagher et al., 2007, Andersen et al., 2018, Morais-Santos et al., 2015), we found that KD of either MCT4 or CD147 nearly ablated the protein expression of the other protein. The chaperone effect was particularly evident for MCT4, where a, presumably misfolded (Wilson et al., 2009), dimer form was predominant when the transporter was overexpressed alone, and this dimer essentially disappeared upon co-expression with CD147. Interestingly and unexpectedly, CD147 overexpression also tripled the mRNA level of MCT4 (but not vice versa). This has to our knowledge not previously been reported and indicates that the interrelationship between the two proteins is more complex than previously anticipated. Although not studied here, the simplest interpretation is that the increased protein expression of CD147 may allow increased MCT4 translation, in turn reducing mRNA degradation. While more difficult to quantify, IFM analysis using line scans, and flow cytometry analysis of cell surface CD147 expression, also indicated that plasma membrane expression of one of the proteins generally increased the plasma membrane expression of the other. MDA-MB-231 cells lack MCT1 due to hypermethylation of its promoter region (Asada et al., 2003), and lactate transport in these cells is therefore mediated by MCT4 (Contreras-Baeza et al., 2019). This allowed us to validate that the reciprocal relationship was also relevant for MCT4 transport function, as shown by increased lactate transport capacity and extracellular lactate accumulation after expression of CD147 or co-expression of both proteins, supporting previous findings in other cancer types (Schneiderhan et al., 2009, Marchiq et al., 2015).

Confirming and extending earlier reports by us and others (Gallagher et al., 2009, Kong et al., 2016, Zhu et al., 2014), we next showed that MDA-MB-231 cell migration and invasion were attenuated by KD of either MCT4 or CD147 and conversely, increased by their overexpression. A role for MCT4 interaction with integrins has been proposed for its role in cell migration based on co-immunoprecipitation and co-localization to the leading edge (Gallagher et al., 2009). However, the mechanisms through which MCT4 favors invasiveness are unknown. MCT4-mediated lactate-H^+^ co-extrusion could, similar to what has been demonstrated for other net acid-extruding transporters, generate a local acidic environment favoring ECM degradation by enhancing protease activity (Stock et al., 2008). Alternatively, or concomitantly with this, the lactate extruded by MCT4 could activate the lactate receptor, HCAR1, further enhancing invasiveness (Ishihara et al., 2022).

We therefore hypothesized that a key role for the MCT4-CD147 co-structure in cancer cell invasiveness could be to stimulate ECM degradation. Consistent with our hypothesis, we first showed that degradation of fluorescent gelatin was decreased by knockdown of MCT4 or CD147 and tended to be increased upon their joint overexpression. MCT4 and CD147 co-localized strongly to protrusive plasma membrane regions that further co-expressed p-cortactin, pSrc, and F-actin. These were also the regions with the most prominent matrix degradation, identifying them as invasive structures, likely invadopodia. Seeding cells on collagen-I, we next demonstrated that overexpression of CD147 and even more so, both MCT4 and CD147, increased the appearance of the reC1M collagen-I neoepitope, which is the cleavage product resulting from collagen-I degradation by MMP-2, 9, or -13 (Leeming et al., 2011). In support, clinically, reC1M measured in serum has been found to be associated with poor survival outcomes in two metastatic breast cancer cohorts (Lipton et al., 2018). MMP2 (Itoh et al., 2001) and MMP13 (Knäuper et al., 2002) are activated by MMP14 (MT1-MMP), a key player in cancer cell invasion and also itself involved in direct degradation of ECM (Poincloux et al., 2009). MMP14 expression was, however, not altered by MCT4 and/or CD147 overexpression, and was in fact mainly localized to large, intracellular vesicle-like structures, although it was also found in the peripheral invasive structures mentioned above. ECM degradation in cancer cells is not only extracellular, and early studies reported its occurrence in unusually large, very acidic vesicles (LAVs) (Montcourrier et al., 1994, Montcourrier et al., 1990). LAVs were reported to be phagosomal structures harboring cathepsin D and multivesicular bodies and of extremely low pH (∼pH 4, compared to ∼pH 5 in lysosomes (Montcourrier et al., 1994, Montcourrier et al., 1990).

We therefore focused our attention on the intracellular structures and found that CD147, MCT4, and MMP14 strongly co-localized in them. They were very large intracellular vesicles, reminiscent of LAVs, while their characteristic F-actin decoration reminded us of the actin comet tails on autophagosomes (Kast et al., 2015, Kast and Dominguez, 2017). Consistent with the latter, MCT4, CD147 and MMP14 co-localized with the lysosomal marker LAMP1 and partially with the autophagosomal marker LC3 in intracellular vesicles, suggesting that a fraction of these vesicles might be destined for autophagic degradation. However, the only partial colocalization with LC3, and the fact that MCT4 and/or CD147 overexpression did not appear to increase total cellular autophagy, seems to argue against a major role of autophagy in MCT4/CD147-facilitated ECM degradation.

Interestingly, MMP14 can be released to the ECM via docking of multivesicular exosomes (MVE) resulting from the fusion of late multivesicular endosomes with the plasma membrane at sites of invadopodia (Hoshino et al., 2013). A hypothesis consistent with our data and the literature is, therefore, that the LAVs are MVE precursors – similar to the endosome/lysosome vesicles which deliver MMPs to invadopodia (Poincloux et al., 2009, Pedersen et al., 2020) - and that these are the MCT4- and CD147-containing vesicles we identify here. MCT4 and CD147 would thus be retrieved from the general plasma membrane along with MMP14 and delivered to invadopodia, consistent with their increased expression in these structures. Consistent with this notion, CD147 and MCT1 were found to be greatly enriched in MVEs released by malignant gliomas (Thakur et al., 2020). Given that MCT1 and MCT4 share their chaperone dependence on CD147, it seems reasonable to suggest that in cells such as MDA-MB-231 cells, which do not contain MCT1, MCT4 would instead be cotransported with CD147 to exosomes. Whether their role in ECM degradation involves a role in formation or recycling of the large vesicles and/or their activity in invadopodia, where increased lactate-H^+^ efflux by MCT4 would create a local, pericellular acidic zone enhancing protease activity, remains to be determined. A working hypothesis is shown in Fig. 6g. It is however interesting in this regard that in gliomas, MCT1 and CD147 were not only enriched in the exosomes but also played a causal role in their release, apparently through raising intracellular free Ca^2+^ (Thakur et al., 2020). It is also feasible that MCT4 activity in these vesicles could contribute to their acidification. Such possibilities, to be studied in future work, not only point to new understanding of the role of MCT4 in cancer, but also have potential for its non-invasive detection in cancer patients for diagnostic- or treatment monitoring purposes.

In conclusion, we show here that MCT4 and CD147 reciprocally regulate each other and support breast cancer cell migration and invasiveness in an interdependent manner. Mechanistically, this involves an MCT4- and CD147-dependent stimulation of ECM degradation and specifically of MMP-mediated collagen-I degradation. MCT4 and CD147 co-localize strongly to invadopodial peripheral structures, as well as in large, LAMP1-positive vesicles, in both cases co-localizing with MMP14, and we suggest that these vesicles mediate delivery of MMP14 to invadopodia.

## Materials and Methods

### Cell lines and cell culture

MDA-MB-231 cells were cultured in Dulbecco’s modified eagle medium (DMEM) 1885 (Substrat og SterilCentralen, #15, University of Copenhagen, Panum 13.01.111) supplemented with 10% Fetal Bovine Serum (FBS, Gibco, #F9665), 1% Penicillin/Streptomycin (Pen/Strep, Sigma-Aldrich, #P0781) and 1% minimum essential medium (MEM) Non-Essential Amino Acids 100x (NEAA, Gibco, #11140-035). For MDA-MB-231 with stable shRNA-mediated MCT4 KD (described in (Andersen et al., 2018)) the media was supplemented with 1 µg/ml puromycin (Gibco, #A11138-02). MCF10A cells were cultured in DMEM containing 4.5 g/l D-glucose, L-glutamine and Pyruvate (Gibco, #41966-029) mixed 1:1 with Nutrient Mixture F-12 (Sigma-Aldrich, #N6658) supplemented with 5% FBS (Gibco, #F9665), 1% Pen/Strep (Sigma-Aldrich, #P0781), 20 ng/mL Epidermal Growth Factor (EGF, Sigma, #E9644), 0.5 µg/mL Hydrocortisone (Sigma-Aldrich, #H0888) and 1% Insulin (Gibco, #41400045). Cells were cultured in culture flasks (T25 or T75: Greiner Bio-one, CELLSTAR, #690160 and #658170) at 37°C, 95% humidity, 5% CO_2_, and passaged when a confluence of 70-80% was reached. Cell cultures were discarded when they reached passage 20-25.

### Knockdown and overexpression

#### Transient knockdown

MDA-MB-231 cells were seeded in 6-well plates (Greiner Bio-One, CELLSTAR, #657-160) and grown to a confluence of ∼25%. On the day of transfection, cells were washed once in transfection medium (MDA-MB-231 culture medium without FBS and Pen/Strep) and 1.5 mL of transfection medium was added to each 6-well. Cells were then treated with 500 µL of transfection medium containing 5 µL Lipofectamine-2000 (Invitrogen, #11668019) and the relevant siRNA, which was applied at final concentrations of 25 nM (CD147 KD) or 50 nM (MCT4 KD). 4-6 h after transfection, 120 µL FBS was added to each 6-well. The medium was replaced with normal growth medium 24 h after transfection, and experiments were performed 48 h after transfection. The following siRNAs, targeting human *SLC16A3* (encoding MCT4) and *BSG* (encoding CD147), all from Dharmacon (via VWR International A/S, Søborg, Denmark), were used (sense strand sequences are given): MCT4 si1: 5’-CCGCAAGGUUACAAGGCAUUU-3’, CD147 si1: 5’-UCCCAGUGCUUGCAAGAUUUU-3’, CD147 si2 (Dharmacon #CTM-465973) sense: 5’-CCACCCACCGCCACAAUAAUU-3’.

#### Overexpression

MDA-MB-231 or MCF10A cells were seeded in 6-well plates (Greiner Bio-One, CELLSTAR, #657-160) and grown to a confluence of 50-70%. Prior to transfection the culture medium in each 6-well was replaced with 2 mL of fresh culture medium. Cells were then treated with 245 µL Opti-MEM (Gibco, #31985062) containing 2.5 µg of the relevant DNA plasmid (for dual expression, 1.25 µg of each plasmid), 7.5 µL Lipofectamine 3000 (Invitrogen, #L3000-015) and 5 µL P3000 Enhancer Reagent (Invitrogen, #L3000-015). After 4-6 h the medium was replaced with fresh culture medium. Experiments were performed 24 h after transfection. Plasmid constructs used were: empty pcDNA3.1 (+) (ctrl.); CD147 in pcDNA3.1 (+), (GenScript, cat #OHu27639); rat MCT4 (rMCT4) in pcDNA3.1(+): rMCT4 in a Xenopus vector (pGHJ-rMCT4) was kindly provided by Holger M. Becker, University of Kaiserslautern, Germany. rMCT4 was subcloned from pGHJ to pcDNA3.1 (+) using restriction enzyme double digestion and sequenced before use.

### SDS page and Western blotting

Western blot analysis was carried out essentially as in (Sjøgaard-Frich et al., 2021). Briefly, cells were grown to ∼80% confluency, washed in ice-cold PBS and lysed (1% SDS, 10 mM Tris-HCl, 1 mM NaVO_3_, cOmplete^TM^ Mini Protease Inhibitor (Roche, #11836153001), pH 7.5, 95°C). Lysates were sonicated, centrifuged, and protein concentrations determined using a Bio-Rad DC Protein assay kit (Bio-Rad, #500-0113, -0114, -0115). Protein concentrations were equalized with ddH_2_O, and lysates were mixed with 1:1 NuPAGE LDS Sample Buffer 4X (Invitrogen, #NP0007) and 0.5 M dithiothreitol (Sigma #646563) and separated by SDS-PAGE using 10% Criterion TGX Precast Midi Protein Gels (BioRad, #567-1035), Tris-Glycine-SDS running buffer (BioRad, #161 0732), and BenchMark protein ladder (Invitrogen, #10747-012). Proteins were transferred to nitrocellulose membranes (BioRad, #170-4159), stained with Ponceau S (Sigma-Aldrich, #7170-1L) and blocked (1 h, 37°C, 5% dry milk in TBST). Membranes were washed 3 x in TBST, incubated with primary antibodies in TBST + 5% BSA (Sigma, #A7906) overnight at 4°C, washed in TBST, incubated with horseradish peroxidase (HRP) conjugated secondary antibodies for 1 h at room temperature (RT), washed in TBST, and developed using Clarity Western ECL Substrate (BioRad, #1705061) on a Fusion FX developer (Vilber). Band intensities were quantified using ImageJ software and normalized to their respective loading control.

### Antibodies used

Primary antibodies against GAPDH (#2118), LC3A/B (#4108), and Golgin-97 (#97537) were purchased from Cell Signaling Technology; against CD147/EMMPRIN (#MAB972) and CD147 (#AF972) from R&D Systems; against MCT4 (#376140) and LAMP-1 (#20011) from Santa Cruz; against MMP14 (#271840 and #192782), p62 (#56416), and p-cortactin (Y421) (#47768) from Abcam; against DCTN1/p150 (#610473), from BD Transduction Laboratories; PDI (#MA3-019) and Rhodamine-conjugated Phalloidin (#R415) were from Invitrogen; against TOM20 (#11802-1-AP) from Proteintech; and against Src (#OP07) from Calbiochem. Secondary antibodies for Western blotting: HRP-conjugated goat-anti-mouse (DAKO, #P0447) and HRP-conjugated goat-anti-rabbit (DAKO, #P0448). All secondary antibodies for immunocytochemistry were from Invitrogen, raised in donkey, and used at 1:600 dilution. anti-goat Alexa Fluor 488 #A11055; anti-goat Alexa Fluor 568 #A11057; anti-goat Alexa Fluor 647 #A21447; anti-mouse Alexa Fluor 568 #A10037; anti-mouse Alexa Fluor 647 #A31571; anti-rabbit Alexa Fluor 488 #A21206; anti-rabbit Alexa Fluor 568 #A10042; anti-rabbit Alexa Fluor 647 #A31573.

### qPCR analysis

Total RNA was isolated using NucleoSpin RNA II (Macherey-Nagel) and reverse transcribed using Superscript III Transcriptase (Invitrogen, #18080044). cDNA was amplified in triplicates utilizing SYBR Green (Roche, #04913914001) in an ABI7900 qPCR machine with the following steps: 95°C for 10 min, 40 cycles of [95°C for 30 s, 55-63°C (depending on the primer pair) for 1 min, 72°C for 30 s], 95°C for 1 min. Primers were designed using NCBI and Primer-BLAST (www.ncbi.nlm.nih.gov), synthesized by Eurofins and diluted in nuclease-free H_2_O (2 µM working dilution). mRNA levels were determined using the Pfaffl method and housekeeping genes β-actin or TATA-binding protein (TBP) and are shown relative to the mRNA level in ctrl. cells. Primers used: CD147, forward: 5’-CCGTAGAAGACCTTGGCTCC-3’, reverse: 5’-TACTCTCCCCACTGGTCGTC-3’. MCT4, forward: 5’-TGCCATTGGTCTCGTGCTG-3’, reverse: 5’-TCTGCCTTCAGGAAGTGCTC-3’. β-actin, forward: 5’-AGCGAGCATCCCCCAAAGTT-3’, reverse: 5’-GGGCACGAAGGCTCATCATT-3’. TBP, forward: 5’-ACCCACCAACAATTTAGTAGTTA-3’, reverse: 5’-GCTCTGACTTTAGCACCTGTTA-3’.

### Immunofluorescence microscopy (IFM) analysis and line scan quantifications

Cells were grown on glass coverslips which, if so specified, were pre-coated with Oregon-green-conjugated gelatin (Invitrogen, #G13186). Cells were washed in ice-cold PBS, fixed in 2% paraformaldehyde (PFA, Sigma, #47608), permeabilized in 0.5% Triton X-100 (Sigma, #X100) in TBST for 5 min, washed 2 x 5 min in TBST, and blocked in 5% BSA (Sigma, #A7906) in TBST for 30 min. Coverslips were incubated for overnight at 4°C with primary antibodies in 1% BSA in TBST, washed 3 x 5 min in 1% BSA in TBST, incubated with Alexa-Fluor-conjugated secondary antibodies for 1 h at RT, and washed in 1% BSA in TBST for 5 x 5 min, with 4’,6-Diamidino-2-Phenylindole (DAPI, Invitrogen, D3571) 1:1000 in 1% BSA included in the second wash. For LAMP1 staining, 0.5% saponin (Sigma, #47036) was additionally present at all steps in the protocol where BSA was used. Coverslips were mounted using 2% N-propyl gallate (Sigma, #P-3130), sealed with nailpolish, and proteins visualized in an Olympus IX83 microscope with a Yokogawa spinning disc, using confocal imaging, a 60X/1.4 NA oil immersion objective, and CellSens Dimension software. Image overlays, intensity adjustments, and maximal intensity z-projections were carried out in ImageJ software. Where indicated, z-stack deconvolution was carried out using CellSens software. Plasma membrane localization was quantified using the ImageJ Color Profiler plugin. All analyses were performed blindly to avoid any bias. For each condition 4-6 images were analyzed. A line selection was drawn across the plasma membrane. From each line scan, the maximum intensity value was used to calculate the mean value for each individual image. Conditions were unblinded, and a mean intensity value was calculated for each condition.

### Flow cytometry

Cells transfected as described above were detached, resuspended in growth medium, spun down at 80 RCF for 3 min and resuspended in flow buffer (1xPBS containing 2% FBS and 0.1% sodium azide). 1 million cells per condition were then stained with phycoerythrin-(PE)-conjugated anti-human CD147 antibody diluted 1:1000 in flow buffer (BioLegend #306211) for 20 min on ice. A sample with 1 million vector-transfected cells incubated in flow buffer without antibody was included as an unstained control sample. Cells were subsequently fixed in fix buffer (PBS containing 2% FBS and 1% PFA) for 20 min on ice, resuspended in flow buffer and stored at 4°C until analyzed on a BD FACS Calibur cytometer, followed by subsequent data analysis using FlowLogic v. 8.6 (Inivai Technologies). The following gating strategy was employed; Cells of interest were selected based on an FSC vs. SSC plot, and a gate marking the CD147-negative population was defined based on the unstained vector ctrl sample. The stained vector ctrl sample was used to define the gates containing the CD147-positive population and the CD147-OE population. Results are shown as % cells in specified gates.

### Analysis of lactate-driven pH_i_ changes

MDA-MB-231 cells were seeded in wilco glass-bottom wells (WillCo-dish, HBST-3522) at a density of 250,000 cells per well 24 h prior to transfection. Cells were transfected as described above, and used for pHi measurements 24 h after transfection. Cells were washed once in Ringer solution (130 mM NaCl, 3 mM KCl, 20 mM HEPES, 1 mM MgCl_2_, 1 mM CaCl_2_, 6 mM NaOH, adjusted to pH 7.4), and incubated with 2 mL Ringer with 4 µM of SNARF^TM^-5F 5-(and-6)-carboxyl acid, AM ester, acetate (Invitrogen, S23923) for 30 min at 37°C (without CO_2_). Cells were washed 3 x in Ringer solution to remove excess SNARF and incubated again for 15 min at 37°C. One mL of Ringer was added to the dish, which was placed on the heated microscope stage (37°C). SNARF fluorescence was measured at 580 and 660 nm after excitation at 488 nm using an epifluorescence microscope (Nikon Eclipse Ti2). Steady state pH_i_ was measured every 5 s for 5 min, after which 1 mL Ringer with 40 mM lactate (Sigma, L7022) was added to reach a final lactate concentration of 20 mM, and image acquisition was re-initiated immediately, with measurements approx. every second for 5 min. For each measurement 11-13 ROIs were chosen and the ratio of the intensity at 580 and 660 nm was calculated after background subtraction. The initial rate of change in pHi was calculated by linear regression in the first 5 time points after adding lactate.

### Lactate concentration measurements

MDA-MB-231 cells were seeded in 60 mm Petri dishes (Greiner Bio-One, CELLSTAR, #628-160) at 520,000 cells per dish, and transfected as described above on the following day. 48 h post transfection the culture medium was collected to measure the lactate concentration. A control sample of unspent culture medium was included to measure background fluorescence. Samples were diluted 1:400 in Milli-Q water and lactate concentration was measured using a Lactate assay kit (Abcam, ab65330) according to the manufacturer’s protocol (using 1.5 µL of Enzyme Mix per well). A set of standards, with lactate concentrations from 0 to 200 pmol were prepared from the reagents provided with the kit. Samples (2.5-3 µL sample adjusted to a final volume of 50 µL per well with Lactate Assay Buffer) were prepared in a black 96-well plate with clear, flat bottom (PerkinElmer, #6005550), and fluorescence was measured at 25°C using a FLUOstar Optima plate reader, Optima software version 1.20-0 and 544/590 nm (ex/em) filters. The Relative fluorescence units (RFU) values of the 0 pmol standard and the culture medium (no cells) sample were subtracted from the RFU value of the samples. The lactate concentration was calculated as: Lactate concentration (mM) = ((La/Sv) x D) x 10-3, where La is the lactate amount calculated from the standard curve (pmol), Sv is the sample volume added in the 96-well (µL) and D is the sample dilution factor. To account for differences in growth rate and cell death during the 48 h of culturing, the lactate concentrations were corrected for cell counts determined at the time of sample collection.

### Transwell migration and invasion assays

Migration and invasion were analyzed using 24-well 8 µm control inserts (Corning, #354578) and matrigel-coated invasion inserts (Corning, #354483), respectively. Before seeding, cells were briefly starved in low serum media (1% FBS, 1% NEAA). In most experiments, 50,000 cells per chamber were seeded onto the inserts in medium containing 1% FBS. Medium containing 10% FBS was added to the lower chambers and chambers were incubated at 37°C and 5% CO_2_ for 20-24 h. After washing with Gurr buffer (Gibco, #10582013), non-migrating/invading cells were gently removed using a cotton swab. The cells located on the lower side of the chamber were fixed in icecold absolute methanol for 1 h, washed 2 x in Gurr buffer and stained with 30% Giemsa stain solution (Sigma-Aldrich, #51811-82-6) for 30 min. Membranes were washed 3 x in Gurr buffer and air-dried. Alternatively, 10.000, 25.000 and 45.000 cells, respectively, were seeded per chamber, respectively, cells were fixed in 4% PFA for 20 min at RT, permeabilized in 0.5% Triton-X100 in TBST for 20 min, and stained with DAPI (1:1000) for 10 min at RT. All other procedures were the same as above. Stained membranes were cut out and placed on a glass slide. Membranes were imaged using a 40X/0.9 NA air immersion objective and bright field illumination on an Olympus IX83 microscope. Cells were counted manually from 10-20 images per membrane.

### Quantification of gelatin degradation

*G*lass coverslips were coated with 60°C preheated Oregon-green conjugated gelatin from pig skin (Invitrogen, #G13186) at 0.5 mg/mL in PBS + 2% sucrose (VWR, #27480.294). Coverslips were placed on a piece of parafilm, 45 µL gelatin was added to a coverslip and transferred to the next coverslip by tilting the coverslip vertically with tweezers. To form a thin uniform layer, excess gelatin was removed using gentle aspiration from a vacuum source in the periphery of the coverslip. The gelatin from the second coverslip was transferred back to the Eppendorf tube kept at 60°C, and the procedure was repeated for the remaining coverslips. Coated coverslips were placed in a 12-well plate protected from light and left to dry for ∼1-1.5 h. For fixation, each well was incubated for 15 min on ice with 1 mL of pre-chilled 0.5% glutaraldehyde (Sigma, #G6257) diluted in PBS, followed by 3 washes in PBS at RT. Each well was incubated with 1 mL of freshly prepared 5 mg/mL Sodium borohydride (Sigma-Aldrich, #452882) in PBS for 3 min followed by 3 washes in PBS. Coverslips were stored in 12-well plates in PBS with 1% Pen/Strep at 4°C until use. Cells were seeded on the gelatin-coated coverslips in medium without Pen/Strep, at 70,000 cells per mL. After 4, 8 or 12 h as indicated, coverslips were washed in PBS, fixed in 4% PFA (Sigma, #47608) in PBS for 10 min at RT, followed by 3 washes in PBS. Cells were stained according to the IFM procedure described above using Rhodamine-conjugated phalloidin (RP, Invitrogen, #R415) to visualize F-actin and DAPI to visualize nuclei. Images were obtained employing an Olympus IX83 microscope. The degraded area fraction was quantified using the AdaptiveThreshold plugin and the Area Fraction measurement from the *Measure* tool in ImageJ. Cell counts were obtained by manually counting the nuclei using the multi-point tool in ImageJ. For each image, the degraded area fraction was divided by the cell count. The relative degradation for a given time point and condition was then calculated as the mean of 6-13 images.

### Collagen-I cleavage analysis

6-wells were coated with collagen-I by adding 1 mL of 50 µg/mL collagen type I (Corning, #354249) in 20 mM acetic acid to each well, followed by 1 h and 20 min incubation at 37°C. Remaining collagen solution was removed and wells were rinsed three times with 1 x PBS heated to 37°C. The plate was left to air dry in the cell bench for 1 h, then wrapped in parafilm and stored at 4°C until cells were seeded. Transfected MDA-MB-231 cells were resuspended in culture medium without P/S and seeded in collagen-coated 6-wells at a density of 350,000 cells per well. After 48 h, 1 mL of culture medium were sampled from each condition and stored at -20°C. reC1M (matrix metalloproteinase generated fragments of type 1 collagen) concentrations were determined using a competitive enzyme-linked immunosorbent assay (ELISA) according to the manufacturer’s instructions (Nordic Bioscience, Herlev, Denmark). In brief, 96-well streptavidin-coated plates were coated with a biotinylated synthetic reC1M peptide and incubated for 30 min at 20°C. A calibrator reC1M peptide or relevant samples were added to designated wells followed by the addition of peroxidase-conjugated specific monoclonal antibodies. The plate was incubated overnight at 4°C before tetramethylbenzinidine was added to each well and the plates were incubated again for 15 min at 20°C. All incubation steps included shaking at 300 rpm and plates were washed five times with wash buffer (20 mM Tris, 50 mM NaCl, pH 7.2) after each incubation. Lastly, the reaction was stopped by adding 0.18 M H_2_SO_4_ and absorbance was measured at 450 nm with 650 nm as reference.

### Data analysis and statistics

Data analysis and illustrations were performed in Microsoft Excel, GraphPad Prism, ImageJ and BioRender. Data is shown as representative images or as means with standard error of the mean (SEM) error bars, and represent at least 3 independent experiments, unless stated otherwise. Statistical analyses were performed in GraphPad Prism version 8.4.1. To test for statistical differences between two groups, a two-tailed t-test was performed. One-way Analysis of variance (ANOVA) with Dunnett’s multiple comparison post-test was used when more than two groups were compared. Experiments where cells were seeded from the same culture flask and transiently transfected prior to the experiment were regarded as paired.

## Acknowledgements

We gratefully acknowledge the expert technical assistance of Tanja Larsen and Mette Flinck. Ana Fuentes contributed with early pilot experiments. We are grateful to Holger Becker, Hannover Medical School, and to Marie Kveiborg, Biotechnology Research Institute Copenhagen (BRIC), for the kind gifts of the MCT4 and CD147 constructs, respectively.

## Author contributions

Conceptualization, SM, EES and SFP; Methodology, SM, EES, JTU, RC, MF; Investigation, SM, EES, MP, JTU, RC, MTB, MF; Visualization, SM and SFP; Writing – Original Draft, SM and SFP; Writing – Review and Editing, SM and SFP, with inputs from all authors; Funding Acquisition, SFP; Resources, JTU, NWI, SFP; Supervision, SFP, NWI. All authors have seen and approved the last version of the manuscript.

## Data availability

All data are available from the authors upon reasonable request.

## References

Albrechtsen, R., Wewer Albrechtsen, N. J., Gnosa, S., Schwarz, J., Dyrskjot, L. & Kveiborg, M. 2019. Identification of ADAM12 as a Novel Basigin Sheddase. Int J Mol Sci, 20.

Ames, S., Andring, J. T., Mckenna, R. & Becker, H. M. 2020. CAIX forms a transport metabolon with monocarboxylate transporters in human breast cancer cells. Oncogene, 39, 1710–1723.

Andersen, A. P., Samsøe-Petersen, J., Oernbo, E. K., Boedtkjer, E., Moreira, J. M. A., Kveiborg, M. & Pedersen, S. F. 2018. The net acid extruders NHE1, NBCn1 and MCT4 promote mammary tumor growth through distinct but overlapping mechanisms. Int J Cancer, 142, 2529–2542.

Artym, V. V., Zhang, Y., Seillier-Moiseiwitsch, F., Yamada, K. M. & Mueller, S. C. 2006. Dynamic interactions of cortactin and membrane type 1 matrix metalloproteinase at invadopodia: defining the stages of invadopodia formation and function. Cancer Res, 66, 3034–43.

Asada, K., Miyamoto, K., Fukutomi, T., Tsuda, H., Yagi, Y., Wakazono, K., Oishi, S., Fukui, H., Sugimura, T. & Ushijima, T. 2003. Reduced expression of GNA11 and silencing of MCT1 in human breast cancers. Oncology, 64, 380–388.

Bai, Y., Huang, W., Ma, L. T., Jiang, J. L. & Chen, Z. N. 2014. Importance of N-glycosylation on CD147 for its biological functions. Int J Mol Sci, 15, 6356–77.

Bello, A. B., Kim, D., Kim, D., Park, H. & Lee, S. H. 2020. Engineering and Functionalization of Gelatin Biomaterials: From Cell Culture to Medical Applications. Tissue Eng Part B Rev, 26, 164–180.

Berditchevski, F., Chang, S., Bodorova, J. & Hemler, M. E. 1997. Generation of monoclonal antibodies to integrin-associated proteins. Evidence that alpha3beta1 complexes with EMMPRIN/basigin/OX47/M6. J Biol Chem, 272, 29174–80.

Busco, G., Cardone, R. A., Greco, M. R., Bellizzi, A., Colella, M., Antelmi, E., Mancini, M. T., Dell’aquila, M. E., Casavola, V., Paradiso, A. & Reshkin, S. J. 2010. NHE1 promotes invadopodial ECM proteolysis through acidification of the peri-invadopodial space. FASEB J, 24, 3903–3915.

Contreras-Baeza, Y., Sandoval, P. Y., Alarcón, R., Galaz, A., Cortés-Molina, F., Alegría, K., Baeza-Lehnert, F., Arce-Molina, R., Guequén, A., Flores, C. A., San Martín, A. & Barros, L. F. 2019. Monocarboxylate transporter 4 (MCT4) is a high affinity transporter capable of exporting lactate in high-lactate microenvironments. J Biol Chem, 294, 20135–20147.

Egawa, N., Koshikawa, N., Tomari, T., Nabeshima, K., Isobe, T. & Seiki, M. 2006. Membrane type 1 matrix metalloproteinase (MT1-MMP/MMP-14) cleaves and releases a 22-kDa extracellular matrix metalloproteinase inducer (EMMPRIN) fragment from tumor cells. J Biol.Chem., 281, 37576–37585.

Gallagher, S. M., Castorino, J. J. & Philp, N. J. 2009. Interaction of monocarboxylate transporter 4 with beta1-integrin and its role in cell migration. Am.J.Physiol Cell Physiol, 296, C414–C421.

Gallagher, S. M., Castorino, J. J., Wang, D. & Philp, N. J. 2007. Monocarboxylate transporter 4 regulates maturation and trafficking of CD147 to the plasma membrane in the metastatic breast cancer cell line MDA-MB-231. Cancer Res., 67, 4182–4189.

Grass, G. D., Bratoeva, M. & Toole, B. P. 2012. Regulation of invadopodia formation and activity by CD147. J Cell Sci, 125, 777–88.

Grass, G. D., Dai, L., Qin, Z., Parsons, C. & Toole, B.P. 2014. CD147: regulator of hyaluronan signaling in invasiveness and chemoresistance. Adv Cancer Res, 123, 351–73.

Grass, G. D. & Toole, B. P. 2015. How, with whom and when: an overview of CD147-mediated regulatory networks influencing matrix metalloproteinase activity. Biosci Rep, 36, e00283.

Halestrap, A. P. 2013. Monocarboxylic acid transport. Compr.Physiol., 3, 1611–1643.

Hoshino, D., Kirkbride, Kellye C., Costello, K., Clark, Emily S., Sinha, S., Grega-Larson, N., Tyska, Matthew J. & Weaver, Alissa M. 2013. Exosome Secretion Is Enhanced by Invadopodia and Drives Invasive Behavior. Cell Reports, 5, 1159–1168.

Huang, W., Luo, W. J., Zhu, P., Tang, J., Yu, X. L., Cui, H. Y., Wang, B., Zhang, Y., Jiang, J. L. & Chen, Z. N. 2013. Modulation of CD147-induced matrix metalloproteinase activity: role of CD147 N-glycosylation. Biochem J, 449, 437–48.

Ishihara, S., Hata, K., Hirose, K., Okui, T., Toyosawa, S., Uzawa, N., Nishimura, R. & Yoneda, T. 2022. The lactate sensor GPR81 regulates glycolysis and tumor growth of breast cancer. Sci Rep, 12, 6261.

Itoh, Y., Takamura, A., Ito, N., Maru, Y., Sato, H., Suenaga, N., Aoki, T. & Seiki, M. 2001. Homophilic complex formation of MT1-MMP facilitates proMMP-2 activation on the cell surface and promotes tumor cell invasion. EMBO J, 20, 4782–4793.

Jeannot, P. & Besson, A. 2020. Cortactin function in invadopodia. Small GTPases, 11, 256–270.

Kast, D. J. & Dominguez, R. 2017. The Cytoskeleton-Autophagy Connection. Curr Biol, 27, R318–R326.

Kast, D. J., Zajac, A. L., Holzbaur, E. L., Ostap, E. M. & Dominguez, R. 2015. WHAMM Directs the Arp2/3 Complex to the ER for Autophagosome Biogenesis through an Actin Comet Tail Mechanism. Curr Biol, 25, 1791–7.

Kirk, P., Wilson, M. C., Heddle, C., Brown, M. H., Barclay, A. N. & Halestrap, A. P. 2000. CD147 is tightly associated with lactate transporters MCT1 and MCT4 and facilitates their cell surface expression. EMBO J, 19, 3896–3904.

Knäuper, V., Bailey, L., Worley, J. R., Soloway, P., Patterson, M. L. & Murphy, G. 2002. Cellular activation of proMMP-13 by MT1-MMP depends on the C-terminal domain of MMP-13. FEBS Lett, 532, 127–30.

Ko, S. Y., Lee, W., Weigert, M., Jonasch, E., Lengyel, E. & Naora, H. 2023. The glycoprotein CD147 defines miRNA-enriched extracellular vesicles that derive from cancer cells. J Extracell Vesicles, 12, e12318.

Kong, S. C., Nohr-Nielsen, A., Zeeberg, K., Reshkin, S. J., Hoffmann, E. K., Novak, I. & Pedersen, S. F. 2016. Monocarboxylate Transporters MCT1 and MCT4 Regulate Migration and Invasion of Pancreatic Ductal Adenocarcinoma Cells. Pancreas, 45, 1036–1047.

Le, F. R., Chiche, J., Marchiq, I., Naiken, T., Ilc, K., Murray, C. M., Critchlow, S. E., Roux, D., Simon, M. P. & Pouyssegur, J. 2011. CD147 subunit of lactate/H+ symporters MCT1 and hypoxia-inducible MCT4 is critical for energetics and growth of glycolytic tumors. Proc.Natl.Acad.Sci.U.S.A, 108, 16663–16668.

Lee, H., Overall, C. M., Mcculloch, C. A. & Sodek, J. 2006. A critical role for the membrane-type 1 matrix metalloproteinase in collagen phagocytosis. Mol Biol Cell, 17, 4812–26.

Leeming, D., He, Y., Veidal, S., Nguyen, Q., Larsen, D., Koizumi, M., Segovia-Silvestre, T., Zhang, C., Zheng, Q., Sun, S., Cao, Y., Barkholt, V., Hägglund, P., Bay-Jensen, A., Qvist, P. & Karsdal, M. 2011. A novel marker for assessment of liver matrix remodeling: an enzyme-linked immunosorbent assay (ELISA) detecting a MMP generated type I collagen neo-epitope (C1M). Biomarkers, 16, 616–28.

Liang, Y., Zhang, H., Song, X. & Yang, Q. 2020. Metastatic heterogeneity of breast cancer: Molecular mechanism and potential therapeutic targets. Seminars in Cancer Biology, 60, 14–27.

Lipton, A., Leitzel, K., Ali, S. M., Polimera, H. V., Nagabhairu, V., Marks, E., Richardson, A. E., Krecko, L., Ali, A., Koestler, W., Esteva, F. J., Leeming, D. J., Karsdal, M. A. & Willumsen, N. 2018. High turnover of extracellular matrix reflected by specific protein fragments measured in serum is associated with poor outcomes in two metastatic breast cancer cohorts. Int J Cancer, 143, 3027–3034.

Lu, P., Takai, K., Weaver, V. M. & Werb, Z. 2011. Extracellular matrix degradation and remodeling in development and disease. Cold Spring Harb.Perspect.Biol, 3.

Marchiq, I., Albrengues, J., Granja, S., Gaggioli, C., Pouysségur, J. & Simon, M. P. 2015. Knock out of the BASIGIN/CD147 chaperone of lactate/H+ symporters disproves its pro-tumour action via extracellular matrix metalloproteases (MMPs) induction. Oncotarget, 6, 24636–48.

Mego, M., Mani, S. A. & Cristofanilli, M. 2010. Molecular mechanisms of metastasis in breast cancer--clinical applications. Nat Rev Clin Oncol, 7, 693–701.

Montcourrier, P., Mangeat, P. H., Salazar, G., Morisset, M., Sahuquet, A. & Rochefort, H. 1990. Cathepsin D in breast cancer cells can digest extracellular matrix in large acidic vesicles. Cancer Res, 50, 6045–54.

Montcourrier, P., Mangeat, P. H., Valembois, C., Salazar, G., Sahuquet, A., Duperray, C. & Rochefort, H. 1994. Characterization of very acidic phagosomes in breast cancer cells and their association with invasion. Journal of Cell Science, 107, 2381–2391.

Monteiro, P., Rosse, C., Castro-Castro, A., Irondelle, M., Lagoutte, E., Paul-Gilloteaux, P., Desnos, C., Formstecher, E., Darchen, F., Perrais, D., Gautreau, A., Hertzog, M. & Chavrier, P. 2013. Endosomal WASH and exocyst complexes control exocytosis of MT1-MMP at invadopodia. J Cell Biol, 203, 1063–79.

Morais-Santos, F., Granja, S., Miranda-Goncalves, V., Moreira, A. H., Queiros, S., Vilaca, J. L., Schmitt, F. C., Longatto-Filho, A., Paredes, J., Baltazar, F. & Pinheiro, C. 2015. Targeting lactate transport suppresses in vivo breast tumour growth. Oncotarget., 6, 19177–19189.

Mylvaganam, S., Freeman, S. A. & Grinstein, S. 2021. The cytoskeleton in phagocytosis and macropinocytosis. Curr Biol, 31, R619–R632.

Nakamura, S. & Yoshimori, T. 2017. New insights into autophagosome-lysosome fusion. J Cell Sci, 130, 1209–1216.

Payen, V. L., Mina, E., Van Hée, V. F., Porporato, P. E. & Sonveaux, P. 2020. Monocarboxylate transporters in cancer. Mol Metab, 33, 48–66.

Pedersen, N. M., Wenzel, E. M., Wang, L., Antoine, S., Chavrier, P., Stenmark, H. & Raiborg, C. 2020. Protrudin-mediated ER-endosome contact sites promote MT1-MMP exocytosis and cell invasion. J Cell Biol, 219.

Pedraz-Cuesta, E., Fredsted, J., Jensen, H. H., Bornebusch, A., Nejsum, L. N., Kragelund, B. B. & Pedersen, S. F. 2016. Prolactin Signaling Stimulates Invasion via Na(+)/H(+) Exchanger NHE1 in T47D Human Breast Cancer Cells. Mol Endocrinol, 30, 693–708.

Poincloux, R., Lizarraga, F. & Chavrier, P. 2009. Matrix invasion by tumour cells: a focus on MT1-MMP trafficking to invadopodia. J Cell Sci, 122, 3015–24.

Runwal, G., Stamatakou, E., Siddiqi, F. H., Puri, C., Zhu, Y. & Rubinsztein, D. C. 2019. LC3-positive structures are prominent in autophagy-deficient cells. Scientific Reports, 9, 10147.

Schneiderhan, W., Scheler, M., Holzmann, K. H., Marx, M., Gschwend, J. E., Bucholz, M., Gress, T. M., Seufferlein, T., Adler, G. & Oswald, F. 2009. CD147 silencing inhibits lactate transport and reduces malignant potential of pancreatic cancer cells in in vivo and in vitro models. Gut, 58, 1391–1398.

Seals, D. F., Azucena, E. F., JR., Pass, I., Tesfay, L., Gordon, R., Woodrow, M., Resau, J. H. & Courtneidge, S. A. 2005. The adaptor protein Tks5/Fish is required for podosome formation and function, and for the protease-driven invasion of cancer cells. Cancer Cell, 7, 155–65.

Sjøgaard-Frich, L. M., Prestel, A., Pedersen, E. S., Severin, M., Kristensen, K. K., Olsen, J. G., Kragelund, B. B. & Pedersen, S. F. 2021. Dynamic Na(+)/H(+) exchanger 1 (NHE1) - calmodulin complexes of varying stoichiometry and structure regulate Ca(2+)-dependent NHE1 activation. Elife, 10.

Stock, C., Cardone, R. A., Busco, G., Krahling, H., Schwab, A. & Reshkin, S. J. 2008. Protons extruded by NHE1: Digestive or glue? Eur J Cell Biol.

Sun, J. & Hemler, M. E. 2001. Regulation of MMP-1 and MMP-2 production through CD147/extracellular matrix metalloproteinase inducer interactions. Cancer Res, 61, 2276–81.

Sung, H., Ferlay, J., Siegel, R. L., Laversanne, M., Soerjomataram, I., Jemal, A. & Bray, F. 2021. Global Cancer Statistics 2020: GLOBOCAN Estimates of Incidence and Mortality Worldwide for 36 Cancers in 185 Countries. CA Cancer J Clin, 71, 209–249.

Tang, W. & Hemler, M. E. 2004. Caveolin-1 regulates matrix metalloproteinases-1 induction and CD147/EMMPRIN cell surface clustering. J Biol Chem, 279, 11112–8.

Taylor, P. M., Woodfield, R. J., Hodgkin, M. N., Pettitt, T. R., Martin, A., Kerr, D. J. & Wakelam, M. J. 2002. Breast cancer cell-derived EMMPRIN stimulates fibroblast MMP2 release through a phospholipase A(2) and 5-lipoxygenase catalyzed pathway. Oncogene, 21, 5765–72.

Thakur, A., Qiu, G., Xu, C., Han, X., Yang, T., Ng, S. P., Chan, K. W. Y., Wu, C. M. L. & Lee, Y. 2020. Label-free sensing of exosomal MCT1 and CD147 for tracking metabolic reprogramming and malignant progression in glioma. Science Advances, 6, eaaz6119.

Weigelt, B., Peterse, J. L. & Van’T Veer, L. J. 2005. Breast cancer metastasis: markers and models. Nat Rev Cancer, 5, 591–602.

Wilson, M. C., Meredith, D., Bunnun, C., Sessions, R. B. & Halestrap, A.P. 2009. Studies on the DIDS-binding site of monocarboxylate transporter 1 suggest a homology model of the open conformation and a plausible translocation cycle. J Biol Chem, 284, 20011–21.

Xin, X., Zeng, X., Gu, H., Li, M., Tan, H., Jin, Z., Hua, T., Shi, R. & Wang, H. 2016. CD147/EMMPRIN overexpression and prognosis in cancer: A systematic review and meta-analysis. Sci Rep, 6, 32804.

Zhu, J., Wu, Y. N., Zhang, W., Zhang, X. M., Ding, X., Li, H. Q., Geng, M., Xie, Z. Q. & Wu, H. M. 2014. Monocarboxylate transporter 4 facilitates cell proliferation and migration and is associated with poor prognosis in oral squamous cell carcinoma patients. PLoS.ONE., 9, e87904.

